# SARM1 depletion rescues NMNAT1 dependent photoreceptor cell death and retinal degeneration

**DOI:** 10.1101/2020.04.30.069385

**Authors:** Yo Sasaki, Hiroki Kakita, Shunsuke Kubota, Abdoulaye Sene, Tae Jun Lee, Norimitsu Ban, Zhenyu Dong, Joseph B. Lin, Sanford L. Boye, Aaron DiAntonio, Shannon E. Boye, Rajendra S. Apte, Jeffrey Milbrandt

## Abstract

Leber congenital amaurosis type 9 is an autosomal recessive retinopathy caused by mutations of the NAD^+^ synthesis enzyme NMNAT1. Despite the ubiquitous expression of NMNAT1, patients do not manifest pathologies other than retinal degeneration. Here we demonstrate that widespread NMNAT1 depletion in adult mice mirrors the human pathology, with selective loss of photoreceptors highlighting the exquisite vulnerability of these cells to NMNAT1 loss. Conditional deletion demonstrates that NMNAT1 is required within the photoreceptor. Mechanistically, loss of NMNAT1 activates the NADase SARM1, the central executioner of axon degeneration, to trigger photoreceptor death and vision loss. Hence, the essential function of NMNAT1 in photoreceptors is to inhibit SARM1, highlighting an unexpected shared mechanism between axonal degeneration and photoreceptor neurodegeneration. These results define a novel SARM1-dependent photoreceptor cell death pathway and identifies SARM1 as a therapeutic candidate for retinopathies.

## Introduction

Leber congenital amaurosis (LCA) is a retinal degenerative disease characterized by childhood onset and severe loss of vision. LCA is the most common cause of blindness in children and about 70% of LCA cases are associated with mutations in genes related to the visual cycle, cGMP production, ciliogenesis, or transcription. Recently, more than thirty mutations in the nuclear NAD^+^ biosynthetic enzyme NMNAT1 were identified in patients with autosomal recessive LCA type 9 ^1-6^. Despite the ubiquitous expression of this key NAD^+^ biosynthesis enzyme, LCA9 patients have no other systemic deficits outside the retina. In many cases, LCA9 associated mutant NMNAT1 proteins retain enzymatic activity and other biochemical functions, but appear to be less stable under conditions associated with cell stress ^7^. While it is clear that NAD^+^ deficiency in the retina is an early feature of retinal degenerative disorders in mice ^8,9^, it is not known which cell types and biological pathways are primarily affected in LCA9.

NMNAT1 plays important roles in diverse retinal functions. Overexpression of NMNAT1 in mouse retinal ganglion cells (RGCs) robustly protects against ischemic and glaucomatous loss of the axon and soma ^10^, while conditional ablation in the developing mouse retina causes severe retinal dystrophy and loss of retinal function ^11,12^. Mice harboring *Nmnat1* mutations (V9M and D243G) exhibit severe retinal degeneration while the most common LCA9 mutation (E257K), which is not fully penetrant ^13^, induces a milder retinal degeneration phenotype ^11,14^. In retinal explants, NMNAT1 promotes the survival of mouse retinal progenitor cells ^15^. The requirement for NMNAT in retina is evolutionarily conserved, as the *Drosophila* NMNAT isoform, dNMNAT, is required for the survival of photoreceptor cells after exposure to intense light ^16,17^.

The selective loss of photoreceptor cells in LCA9 suggests the survival and function of these cells are extremely sensitive to deranged NAD^+^ metabolism. Indeed, many of the enzymes involved in photoreceptor function are dependent on NAD^+^ as a cofactor, and for some of these proteins mutations in their corresponding genes lead to blindness. These include variants in the NAD^+^ or NADPH dependent retinal dehydrogenases like RDH12 that cause LCA13 ^18^ and the GTP synthesis enzyme IMPDH1 that causes retinitis pigmentosa ^19,20^. SIRT3, the mitochondrial NAD^+^-dependent deacetylase is also important for photoreceptor homeostasis ^9,21^. Together, these observations highlight the importance of cytosolic NAD^+^ dependent pathways in retinal function ^9,22^; however, the molecular roles of nuclear NAD^+^ and NMNAT1 in the retina are largely unknown.

Multiple enzymatic pathways utilizing distinct metabolic precursors participate in NAD^+^ biosynthesis ^23^. However, in each case, these pathways converge at an NMNAT-dependent step that generates either NAD^+^ or its deamidated form NaAD from the precursor NMN or NaMN. Among the three mammalian NMNAT isoforms, NMNAT1 is the only enzyme localized to the nucleus ^24^. However, in photoreceptors NMNAT1 is present in photoreceptor outer segments ^25^, consistent with an additional, extra-nuclear role of NMNAT1 in photoreceptor cells. This is of particular interest because engineered non-nuclear variants of enzymatically-active NMNAT1 can potently inhibit pathological axon degeneration, which is commonly observed in the early stages of many neurodegenerative disorders ^26-28^. When NMNAT1 is present in the axon, it can compensate for the injury-induced rapid loss of NMNAT2, the endogenous axonal NMNAT ^29^. NMNAT2 in turn, inhibits SARM1, an inducible NAD^+^ cleavage enzyme (NADase) that is the central executioner of axon degeneration ^29-33^. Hence, mutations in NMNAT1 may promote retinal degeneration through the direct impact on NAD^+^ biosynthesis and/or through the regulation of the SARM1-dependent degenerative program.

In this study, we determined the cell types and molecular mechanisms that cause retinal degeneration in LCA9. Using NMNAT1 conditional mutant mice, we showed that photoreceptors degenerate rapidly after the loss of NMNAT1 and that depletion of NMNAT1 in rod or cone cells is necessary and sufficient for the retinal degeneration. The AAV mediated gene replacement of NMNAT1 in photoreceptors partially rescues the visual impairment caused by loss of NMNAT1. Finally, we determined the mechanism by which loss of NMNAT1 leads to photoreceptor degeneration. Loss of NMNAT1 leads to activation of SARM1 in photoreceptors, much as loss of NMNAT2 leads to SARM1 activation in axons ^34^. Moreover, photoreceptor degeneration is mediated by SARM1 in the absence of NMNAT1, much as axon degeneration and perinatal lethality is mediated by SARM1 in the absence of NMNAT2 ^29,32^. Hence, photoreceptor neurodegeneration in LCA9 shares a deep mechanistic similarity to the pathological axon degeneration pathway. Since the SARM1 pathway is likely druggable ^35,36^, these findings provide a framework for developing new therapeutic strategies for treating patients with LCA9 and potentially other retinal disorders.

## Results

NMNAT1 is a nuclear enzyme that synthesizes NAD^+^, an essential metabolite that is central to all aspects of cellular metabolism. NMNAT1 is indispensable for mouse development ^37^ and recent studies identified causative mutations in *NMNAT1* in patients with Leber congenital amaurosis type 9 (LCA9), a disorder associated with severe, early-onset retinal degeneration and vision loss ^1-6^. Patients with LCA9 have no systemic involvement outside the eye, suggesting that certain cells within the retina are particularly vulnerable to the loss of NMNAT1 function. Since no specific antibodies exist for immunocytochemical analysis of NMNAT1 localization, we determined its expression pattern in the retina using mice expressing an NMNAT1-lacZ fusion protein without the nuclear localization signal. Mice heterozygous for this mutant allele were viable and were used to map NMNAT1 expression by staining retinal sections with X-gal. LacZ staining was detected in the retinal pigment epithelium (RPE), photoreceptor outer segments (OS), inner segments (IS), outer nuclear layer (ONL), outer plexiform layer (OPL), inner nuclear layer (INL), inner plexiform layer (IPL), and ganglion cell layer (GCL) suggesting the ubiquitous expression of NMNAT1 in retina (Supplemental Figure 1A).

LCA9 patients are mutant for NMNAT1 throughout the body, yet their defects are limited to the eye. In an effort to model this, we generated a global knockout using *Nmnat1*^*fl/fl*^:*ActCre*^*ERT2*^ mice harboring homozygous *Nmnat1* floxed alleles (*Nmnat1*^*fl*^) and *ActCre*^*ERT2*^, which expresses a tamoxifen-activated Cre recombinase from the ubiquitous actin promoter. We chose a conditional approach because NMNAT1 knockout embryos are lethal ^37^. We treated 2-month-old *Nmnat1*^*fl/fl*^:*ActCre*^*ERT2*^ and control mice with tamoxifen. We first used RT-PCR to measure *Nmnat1* mRNA in the retina at 21 days after tamoxifen and found that it was significantly decreased in NMNAT1 cKO (*Nmnat1*^*fl/fl*^:*ActCre*^*ERT2*^ + tamoxifen) compared with wild type (WT) mice (Supplemental Figure 1B). To investigate the metabolic consequence of NMNAT1 deletion, we measured the levels of NMN, the substrate for NMNAT1, and NAD^+^, the product of NMNAT1, in the retina at 25 days after tamoxifen injection. There is a significant increase in levels of NMN, presumably because it cannot be consumed by NMNAT1. There is also a mild decrease in NAD^+^ in NMNAT1 cKO mice, although this is not statistically significant, suggesting that other NMNAT enzymes are an additional source of NAD^+^ (Figure 1A, B). We next evaluated retinal pathology at 4 weeks after *Nmnat1* excision using biomicroscopy. Fundus images showed abnormalities including attenuation of blood vessels (Figure 1C, D arrowhead) and the appearance of a honeycomb structure, suggesting exposure of retinal pigment epithelium (RPE) cells (Figure 1C, D arrow) in the mutant animals. Histopathological examination of the retina with hematoxylin and eosin (HE) stained sections showed severe retinal degeneration as evidenced by the reduction of the retina thickness and the thinning of the outer nuclear layer (ONL) at 4 weeks post tamoxifen treatment (Figure 1E, F). Quantitative analysis demonstrated a significant reduction of retinal thickness, especially of the ONL (Figure 1G, H). Hence, photoreceptor cells are highly vulnerable following the loss of NMNAT1.

**Figure 1.**
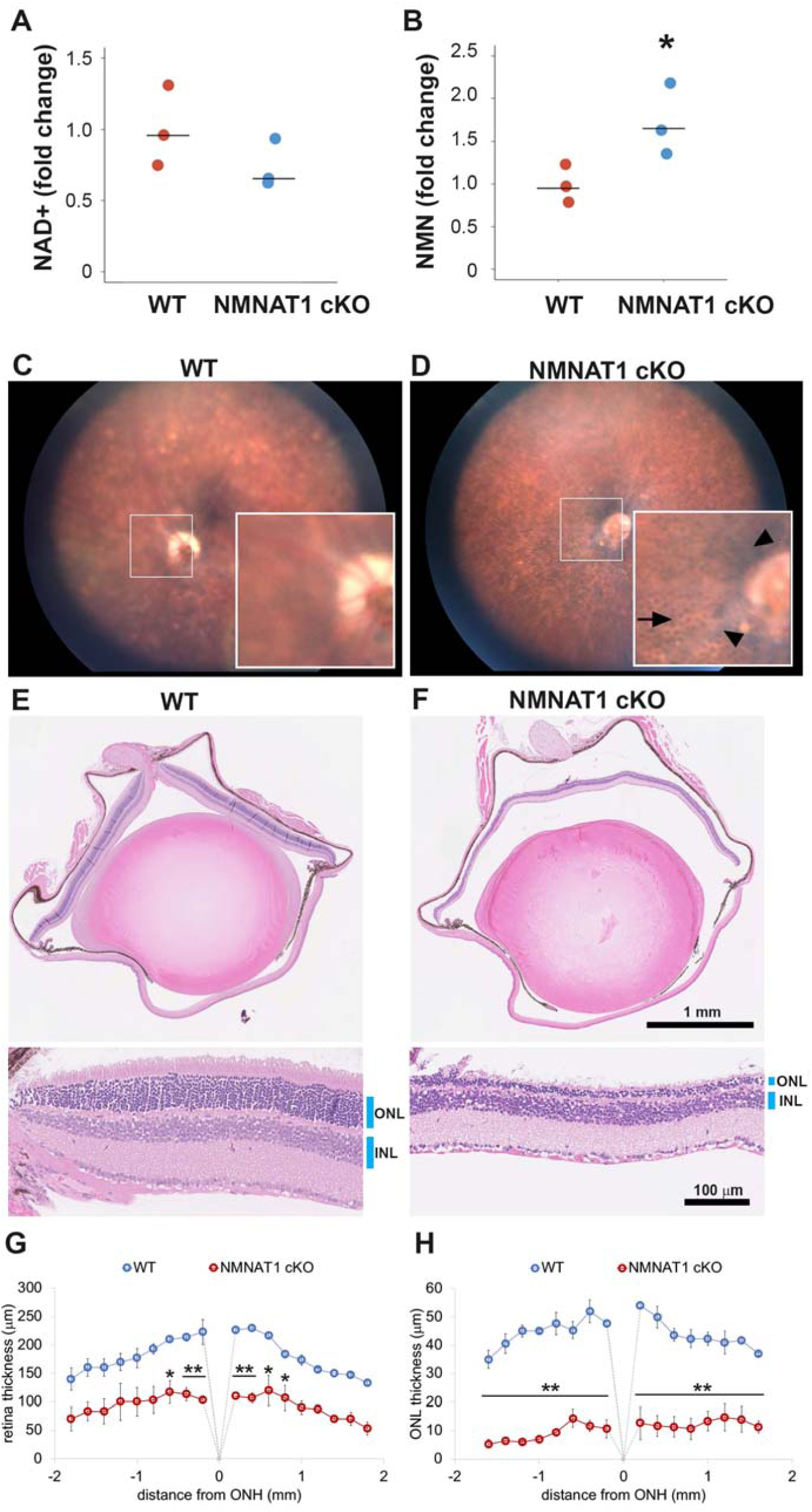
NMNAT1 depletion induces severe retinal degeneration. (A, B) Metabolite analysis by LC-MSMS in retinal tissues from WT or NMNAT1 conditional knockout (*Nmnat1*^*fl/fl*^:*ActCre*^*ERT2*^ + tamoxifen : NMNAT1 cKO) mice at 25 days post tamoxifen injection. Fold changes of NAD^+^ (A) and NMN (B) concentrations compared with that of WT retinal tissues are shown. *p < 0.05 denotes the significant difference from WT with Kruskal-Wallis test (n = 3 mice for WT and n = 3 mice for NMNAT1 cKO). Graphs show the all data points and median (cross bars). (C, D) Fundus biomicroscopy images of the retina from wild type (WT, C) or NMNAT1 conditional knock out (*Nmnat1*^*fl/fl*^:*ActCre*^*ERT2*^ + tamoxifen: NMNAT1 cKO, D) mice at 4 weeks post tamoxifen injection. (E, F) representative images of hematoxylin and eosin stained eye sections from WT mice (E) or NMNAT1 conditional knockout (*Nmnat1*^*fl/fl*^:*ActCre*^*ERT2*^ + tamoxifen: NMNAT1 cKO, E) mice at 4 weeks post tamoxifen injection (ONL: outer nuclear layer and INL: inner nuclear layer). The substantial thinning of the ONL was observed in 3 WT and 3 NMNAT1 cKO mice. (G) The quantification of the retina thickness from WT and NMNAT1 conditional knockout (*Nmnat1*^*fl/fl*^:*ActCre*^*ERT2*^ + tamoxifen: NMNAT1 cKO) mice were shown. Graphs show the average and error bars represent the standard error. Statistical analysis was performed by two-way ANOVA with Tukey post-hoc test (n = 3 mice for WT, n = 3 mice for NMNAT1 cKO (*Nmnat1*^*fl/fl*^:*ActCre*^*ERT2*^ + tamoxifen at 4 weeks post tamoxifen injection)). F(1, 72) = 309, p < 1.0×10^−16^ between WT and NMNAT1 cKO retina. * p<0.05 and ** p<0.001 denotes the significant difference compared with WT retina. (H) The quantification of the ONL thickness from WT and NMNAT1 conditional knockout (*Nmnat1*^*fl/fl*^:*ActCre*^*ERT2*^ + tamoxifen: NMNAT1 cKO) mice were shown. Graphs show the average and error bars represent the standard error. Statistical analysis was performed by two-way ANOVA with Tukey post-hoc test (n=3 mice for WT and n=3 mice for NMNAT1 cKO). F(1, 72) = 1023, p < 1.0×10^−16^ between WT and NMNAT1 cKO retina. ** p<0.001 denotes the significant difference compared WT.

To gain insights into the temporal aspects of the retinal degenerative process, we analyzed retinal morphology at seven time points after tamoxifen administration. The loss of nuclei in the ONL layers were evident at 25 days post tamoxifen injection and robust retinal thinning was evident at 33 days post tamoxifen injection (Figure 2A). We measured the loss of photoreceptor cells by counting the number of ONL cell nuclei. Cell loss was first detected in the ONL around 3 weeks after tamoxifen administration and gradually increased such that only ∼15% of the cells remained at 33 d (Figure 2B). Next, we evaluated retinal function after NMNAT1 deletion using electroretinogram (ERG). We examined three cohorts of mice: *Nmnat1*^*fl/fl*^:*ActCre*^*ERT2*^ treated with tamoxifen, untreated *Nmnat1*^*fl/fl*^:*ActCre*^*ERT2*^ or *Nmnat1*^*fl/fl*^ treated with tamoxifen. In mutant animals in which NMNAT1 was excised, we observed a complete loss of both scotopic (rod-driven responses) and photopic (cone-driven responses) responses, indicating the loss of Nmnat1 in mature retina causes severe photoreceptor dysfunction (Figure 2C-E). This is consistent with previous reports showing developmental retinal defects in the tissue specific Nmnat1 knockout mice ^11,12^. While previous reports show that Nmnat1 is necessary for appropriate retinal development, our pathological and functional analyses of conditional deletion of NMNAT1 in two-month-old mice demonstrates that NMNAT1 is also necessary for photoreceptor cell maintenance and mature retinal functions.

**Figure 2.**
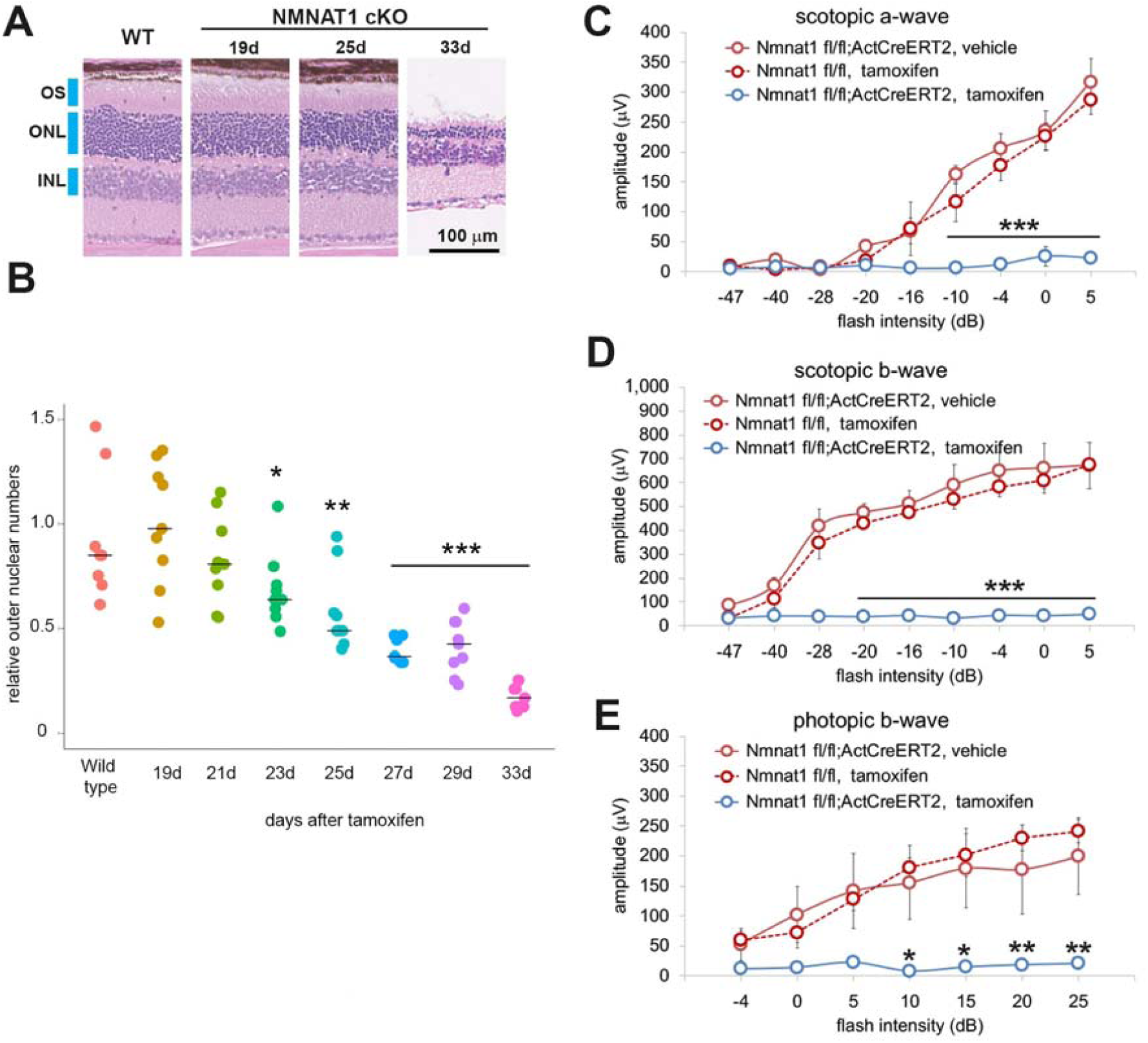
NMNAT1 induces the loss of photoreceptor cells and retinal function. (A) Representative images of hematoxylin/eosin stained sections showing time course of retinal degeneration in NMNAT1 conditional knockout (*Nmnat1*^*fl/fl*^:*ActCre*^*ERT2*^ + tamoxifen: NMNAT1 cKO) mice at 19 to 33 days post tamoxifen injection or littermate wild type mice at 33 days post tamoxifen injection (WT). Blue bars indicate outer nuclear layer (ONL), inner nuclear layer (INL), and outer segment (OS). Similar results were obtained from 3 mice at each time point. (B) Quantification of relative ONL nuclei numbers of NMNAT1 conditional knockout mouse (*Nmnat1*^*fl/fl*^:*ActCre*^*ERT2*^ + tamoxifen: NMNAT1 cKO) compared with WT at various time after tamoxifen injection. The graph shows all data points and median (cross bars). Statistical analysis was performed by one-way ANOVA with Holm-Bonferroni multiple comparison (n = 3 mice for each of WT, 19d, 21d, 33d and n = 4 mice for each of 25d, 27d). F(7, 64) = 19, p = 1.9×10^−13^. * p < 0.05, **p < 0.001, and ***p < 0.0001 denotes the significant difference compared with wild type (WT). (C, D, E) ERG analysis of controls (*Nmnat1*^*fl/fl*^:*ActCre*^*ERT2*^ vehicle or *Nmnat1*^*fl/*fl^ + tamoxifen) and NMNAT1 conditional knockout (*Nmnat1*^*fl/fl*^:*ActCre*^*ERT2*^ + tamoxifen: NMNAT1 cKO). Graphs show the average and error bars represent the standard error. Scotopic a-wave (C), scotopic b-wave (D), and photopic b-wave (E) are shown. Statistical analysis was performed by two-way ANOVA with Tukey post-hoc test (n = 3 mice for *Nmnat1*^*fl/fl*^:*ActCre*^*ERT2*^ with vehicle, n = 3 mice for *Nmnat1*^*fl/fl*^ at 33 days post tamoxifen injection, n = 4 mice for *Nmnat1*^*fl/fl*^:*ActCre*^*ERT2*^ at 33 days post tamoxifen injection). F(1, 72) = 220, p < 2 × 10^−16^ between controls (*Nmnat1*^*fl/fl*^:*ActCre*^*ERT2*^ with vehicle and *Nmnat1*^*fl/fl*^ 33 days post tamoxifen injection) and NMNAT1 cKO for scotopic a wave, F(1, 72) = 633, p < 2 × 10^−16^ between controls and NMNAT1 cKO for scotopic b wave, F(1, 56) = 94, p = 1.3 × 10^−13^ between controls and NMNAT1 cKO for photopic b wave. * p <0.05, **p < 0.001, and ***p < 0.0001 denote a significant difference compared with WT.

In addition to NMNAT1, mammalian cells encode two other NMNAT isoforms; NMNAT2 that is localized in the Golgi and cytosol, and NMNAT3 that is localized inside the mitochondria. Since the loss of NMNAT1 induced retinal degeneration, we wished to determine the role of NMNAT2 and 3 in the retinal structure/function. A previous study showed that NMNAT2 knockout mice are perinatally lethal and have truncated optic nerves as well as peripheral axon degeneration ^38^. We could not assess the role of NMNAT2 in retinal function due to the lack of conditional knockout mice. On the other hand, NMNAT3 deficient mice (Nmnat3^*KO*^) are viable with splenomegaly and hemolytic anemia ^39^. We generated Nmnat3^*KO*^ mice and investigated their retinal function using ERG. Consistent with the previous report, NMNAT3^*KO*^ mice showed splenomegaly (data not shown), however, there were no defects in ERG (Supplemental Figure2). These results indicate that NMNAT3 is dispensable for retinal function, suggesting NMNAT1 is the functionally dominant isoform controlling retinal phenotype.

Identifying the cells that are vulnerable to NMNAT1 loss is key to understanding LCA9 pathogenesis. The severe loss of the ONL nuclei (Figures 2B) induced by NMNAT1 deletion prompted us to test whether loss of NMNAT1 specifically in photoreceptors would result in their death and recapitulate the phenotype observed using the widely expressed *ActCre*^*ERT2*^. We therefore generated mice lacking NMNAT1 specifically in rod photoreceptors (NMNAT1 Rho-Cre) by crossing the *Nmnat1*^*fl/fl*^ mice with Rhodopsin-iCre75 mice in which Cre recombinase expression is driven by the rhodopsin promoter starting postnatally at P7 ^40^. We analyzed the retinas of *Nmnat1*^*fl/fl*^:Rhodopsin-Cre mice at 6-weeks-of-age. Similar to previous findings using Crx-Cre that expresses Cre recombinase in developing photoreceptors as early as E11 ^11,12^, histological analysis revealed severe thinning of the ONL in these mutant mice (Figure 3A, B). The quantitative analysis showed a significant reduction of the retina and ONL thickness in NMNAT1 Rho-Cre retina (Figure 3D) as well as a significant reduction in ONL cell number as detected by nuclear counts (Figure 3F). Consistent with the loss of ONL cells, ERG analysis showed a severe reduction in the scotopic-a and –b waves, representing rod photoreceptor function, in the *Nmnat1*^*fl/fl*^:RhodopsinCre mice (Figure 3G, H). In addition, we found decreases in cone mediated photoresponses (photopic b-wave signal) (Figure 3I) that is likely secondary to a loss of rod photoreceptor cells due to loss of required rod-derived survival factors ^9,41^.

**Figure 3.**
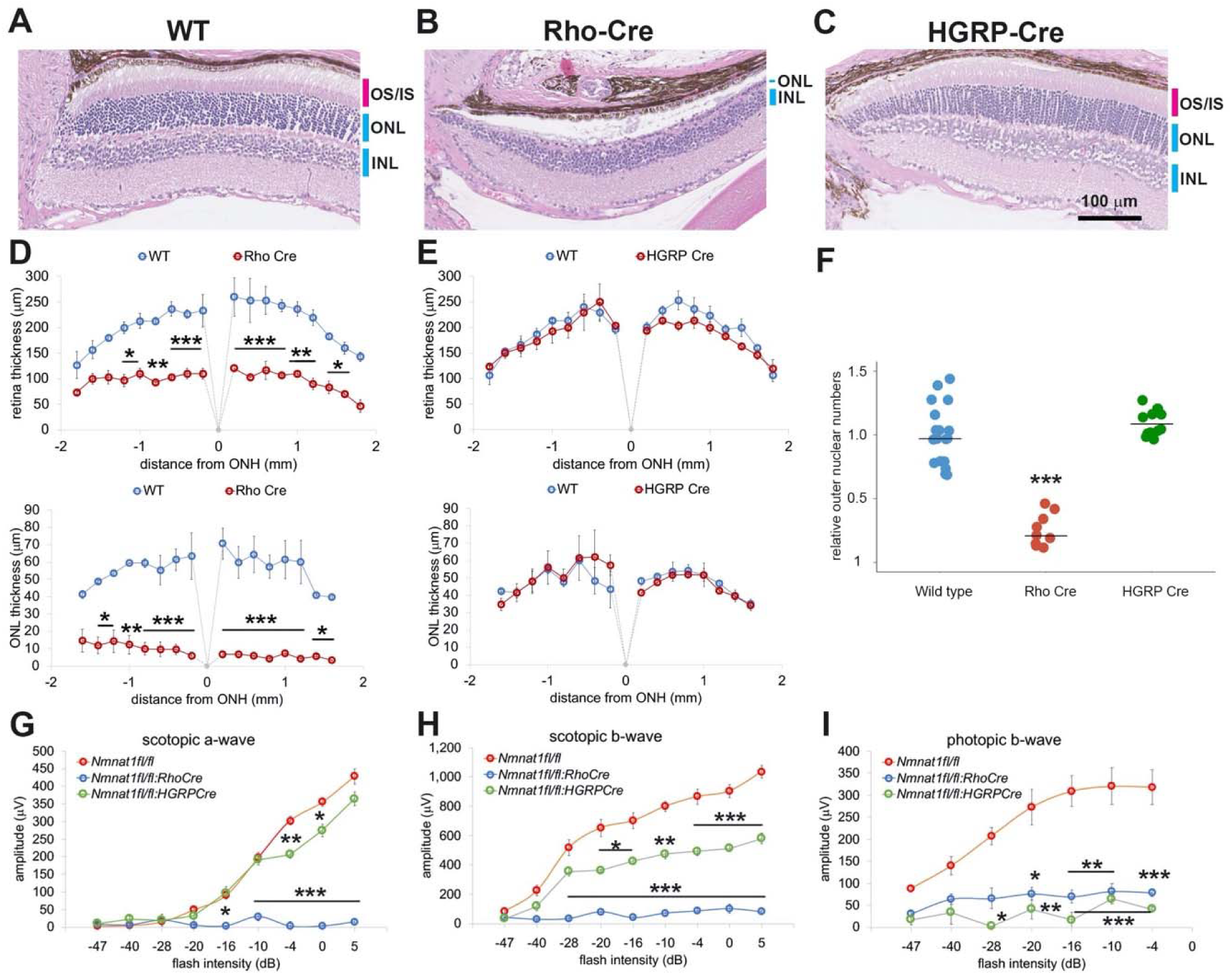
Photoreceptor specific depletion of NMNAT1 induces retinal degeneration. (A, B, C) Hematoxylin and eosin stained eye sections from 6-week old wild type (WT, A), rod-specific NMNAT1 KO (*Nmnat1*^*fl/fl*^:*Rho-Cre*: Rho-Cre, B), or cone-specific NMNAT1 KO (*Nmnat1*^*fl/fl*^:*HGRP-Cre* : HGRP-Cre, C) mice. Blue bars indicate outer nuclear layer (ONL) and inner nuclear layer (INL). Red bars indicate the outer segments (OS) and inner segments (IS). Similar results were obtained from 3 mice for each genotype. (D) Quantification of retina and ONL thickness in WT or rod-specific NMNAT1 KO (Rho-Cre) retinas. The graph shows all data points and median (cross bars). Statistical analysis was performed by two-way ANOVA with Tukey post-hoc test (n = 3 mice for WT and n = 3 mice for Rho-Cre). F(1, 72) = 428, p < 2 × 10^−16^ between WT and Rho-Cre retina thickness and F(1, 64) = 530, p < 2 × 10^−16^ between WT and Rho-Cre ONL thickness. * p < 0.05, **p < 0.001, and ***p < 0.0001 denote significant differences compared with WT. (E) Quantification of retina and ONL thickness in WT or cone-specific NMNAT1 KO (HGRP-Cre) retinas. Graphs show the average and error bars represent the standard error. Statistical analysis was performed by two-way ANOVA with Tukey post-hoc test (n = 3 mice for WT and n = 3 mice for HGRP-Cre). F(1, 72) = 4, p = 0.037 between WT and HGRP-Cre retina thickness and F(1, 64) = 0.03, p = 0.87 between WT and HGRP-Cre ONL thickness. There are no significant differences in HGRP retina and ONL thickness compared with WT. (F) Quantification of relative ONL nuclei numbers compared with WT. The graph shows the all data points and median (cross bars). Statistical analysis was performed by one-way ANOVA with Holm-Bonferroni multiple comparison (n = 6 mice for WT, n = 3 mice for Rho-Cre, n = 3 mice for HGRP-Cre). F(2, 35) = 59, p = 5.9×10^−12^. ***p < 0.0001 denotes the significant difference compared WT. (G, H, I) ERG analysis of WT, Rho-Cre, and HGRP-Cre mice. Scotopic a-wave (G), scotopic b-wave (H), and photopic b-wave (I) are shown. Graphs show the average and error bars represent the standard error. Statistical analysis was performed by two-way ANOVA with Tukey post-hoc test (n = 6 mice for WT, n = 3 mice for Rho-Cre, n = 3 mice for HGRP-Cre). F(2, 81) = 314, p < 2.0 × 10^−16^ among genotypes (WT, Rho-Cre, and HGRP-Cre) for scotopic a wave, F(2, 81) = 413, p < 2 × 10^−16^ among genotypes for scotopic b wave, F(2, 63) = 102, p < 2 × 10^−16^ among genotypes for photopic b wave. * p <0.05, **p < 0.001, and ***p < 0.0001 denote a significant difference compared with WT.

To explore directly the role of NMNAT1 in cones, we deleted NMNAT1 using the cone-specific HGRP-Cre. We crossed *Nmnat1*^*fl/fl*^ mice with HGRP-Cre mice in which Cre recombinase expression is driven by the human red/green pigment promoter starting at P10 ^42^. At 6-weeks-of-age we examined these mutant mice histologically, but did not detect any gross abnormalities, presumably due to the low number of cones (only 3% of total photoreceptors) in mice (Figure 3A, C, E). However, ERG analysis showed a complete loss of the photopic-b wave, which is derived from cone photoreceptors. This functional result demonstrates that NMNAT1 activity is vital for cone function (Figure 3I). In summary, these genetic ablation experiments demonstrate the importance of NMNAT1 for proper function and survival of both rods and cones, and indicate that LCA9-associated retinal degeneration is likely due to the direct cell-autonomous effects of *NMNAT1* mutations in photoreceptors.

The loss-of-function studies above demonstrate that NMNAT1 is necessary in photoreceptors for proper retinal function. Next we assessed whether viral-mediated expression of NMNAT1 in photoreceptors in an otherwise NMNAT1 deficient animal is sufficient to promote retinal function. First, we developed a system for retinal expression of transgenes. We subretinally delivered AAV8(Y733F) containing the photoreceptor specific human rhodopsin kinase (hGRK1) promoter driving GFP ^43,44^. Virus was injected into the subretinal space of two-month-old wildtype mice. Transgene expression was evaluated 4-6 weeks post-injection. AAV-mediated GFP expression was observed in a subset of rhodopsin-positive cells but was weak in the inner nuclear layer (INL) (Supplemental Figure 3A). These results confirm earlier reports that the hGRK1 promoter restricts transgene expression primarily to photoreceptors ^43^. We next asked whether AAV-mediated expression of HA-tagged human NMNAT1 could prevent retinal degeneration caused by NMNAT1 excision. Two-month-old Nmnat1^*fl/fl*^;*ActCre*^*ERT2*^ *mice* received subretinal injections of AAV-NMNAT1 in one eye, and control vector (AAV-GFP) in the contralateral eye. We confirmed the expression of NMNAT1-HA in a subset of outer nuclear cells and a minor population of inner nuclear cells (Supplemental Figure 3B, C). Mice that received AAV-NMNAT1 and AAV-GFP were then treated with tamoxifen to deplete endogenous NMNAT1. One month after tamoxifen treatment, we examined retinal function. Despite the expression of NMNAT1 in only a subset of photoreceptors, we observed significantly increased scotopic a-wave amplitudes in AAV-NMNAT1 treated retinas compared with retinas injected with AAV-GFP (Supplemental Figure 3D) accompanied by a small increase in the scotopic and photopic b-wave amplitudes between AAV-NMNAT1 and AAV-GFP treated retinas (Supplemental Figure 3E, F). Hence, NMNAT1 gene delivery to photoreceptor cells significantly improved their function in this LCA9 model.

We next sought to determine the molecular mechanisms required for retinal degeneration in the NMNAT1-deficient retina. In injured peripheral nerves, the loss of NMNAT2 induces an increase in NMN that is hypothesized to activate SARM1-dependent axon degeneration ^45,46^. Our metabolomic analysis revealed that NMN is increased in the NMNAT1-deficient retinas (Figure 1B), and previous studies have detected SARM1 in mouse and bovine photoreceptor cells ^25,47,48^. These results raised the possibility that the increased retinal NMN activates SARM1 NADase activity, inducing NAD^+^ loss and cellular degeneration in the retina. To test this hypothesis, we crossed *Nmnat1*^*fl/fl*^:*ActCre*^*ERT2*^ mice with SARM1 knockout mice ^49^ to generate *Nmnat1*^*fl/fl*^:*ActCre*^*ERT2*^:*Sarm1*^*KO*^ mice. NMNAT1 was excised in these mice via tamoxifen administration at 2 months of age (NMNAT1 cKO: SARM1 KO). First, we assessed SARM1 activation via measurement of cADPR, a product of the SARM1 NAD+ cleavage enzyme and a biomarker of SARM1 activity as well as NAD+ ^34^. While the loss of NAD+ was not statistically significant at 25 days post tamoxifen injection in NMNAT1 cKO retina (Figure 1A), there was significant loss of NAD+ at 29 to 32 days post tamoxifen in NMNAT1 cKO but not in NMNAT1cKO:SARM1 KO retina (Figure 4A) These data suggest activation of the SARM1 NADase in NMNAT1-deficient retina. Consistent with this idea, we also observed a significant increase of cADPR in NMNAT1 cKO retina in a SARM1 dependent manner (Figure 4B). Hence, SARM1 is activated by the loss of NMNAT1. Next, we assessed retinal degeneration. While there is a dramatic loss of ONL nuclei 32 d post tamoxifen injection in NMNAT1 cKO retina, there was no obvious loss of ONL cells in NMNAT1 cKO: SARM1 KO retina (Figure 4C-E). Quantitative analysis showed no reduction of retinal and ONL thickness in NMNAT1 cKO:SARM1 KO retina compared with WT (Figure 4F, G) in sharp contrast to the dramatic loss of retinal and ONL thickness in the NMNAT1 cKO (Figures 1 and 2). Moreover, there was no detectable loss of ONL nuclei in NMNAT1 cKO:SARM1 KO compared with WT, demonstrating that SARM1 is necessary for photoreceptor cell death induced by the loss of NMNAT1 (Figure 4H). We next examined the functional consequences of NMNAT1 depletion in the presence or absence of SARM1 using ERGs, and again found that loss of SARM1 prevented the severe loss of both scotopic and photopic responses due to NMNAT1-deficiency (Figure 4I-K). Taken together, these findings demonstrate that loss of NMNAT1 leads to the activation of SARM1, and that SARM1 is required for the subsequent photoreceptor degeneration and loss of visual function. Therefore, the essential function of NMNAT1 in photoreceptors is to inhibit SARM1, and inhibition of SARM1 is a candidate therapeutic strategy for the treatment of LCA9.

**Figure 4.**
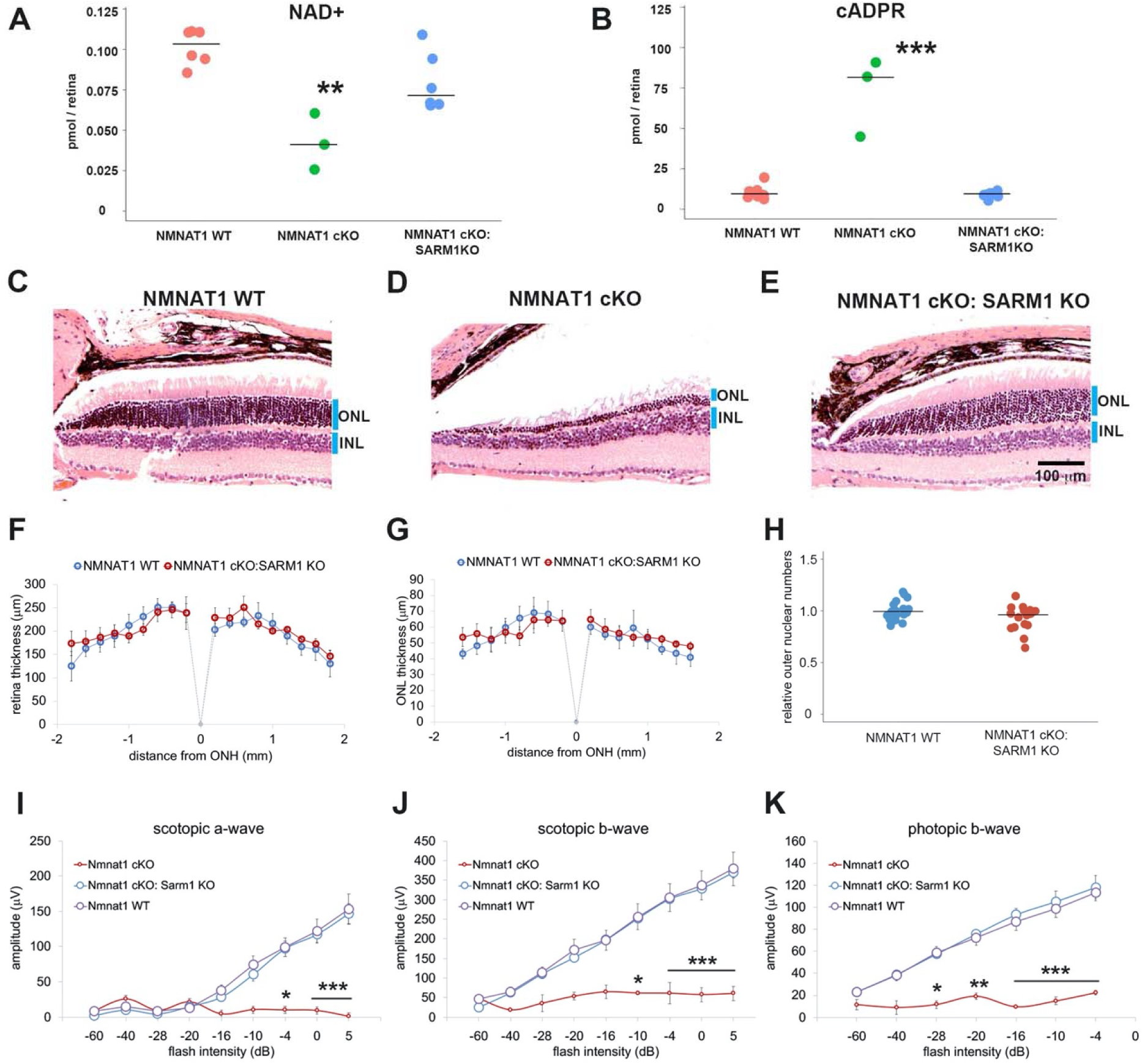
Depletion of SARM1 rescues retinal degeneration in the NMNAT1-deficient retina. (A, B) Metabolite analysis by LC-MSMS in retinal tissues from WT, NMNAT1 conditional knockout (*Nmnat1*^*fl/fl*^:*ActCre*^*ERT2*^ + tamoxifen at 29 to 32 days post tamoxifen injection: NMNAT1 cKO), or NMNAT1 cKO:SARM1 KO mice were shown. Metabolites form whole retina of one eye were analyzed for NAD^+^ (A) and cADPR (B). concentrations compared with that of WT retinal tissues are shown. Graphs show the all data points and median (cross bars). Statistical analysis was performed by one-way ANOVA with Holm-Bonferroni multiple comparison (n = 7 mice for WT, n = 3 mice for NMNAT1 cKO, and n = 6 mice for NMNAT1 cKO:SARM1 KO). F(2, 13) = 259, p = 3.0×10^−4^. For NAD+ and F(2,13) = 48, p = 9.43 × 10^−7^ for cADPR. **p < 0.001 and ***p<0.0001 denotes the significant difference compared WT. (C, D, E) Representative images of hematoxylin and eosin stained eye sections from NMNAT1 WT (*Nmnat1*^*fl/fl*^: *Sarm1*^*+/-*^, C), NMNAT1 cKO (*Nmnat1*^*fl/fl*^:*Sarm1*^*+/+*^: *ActCre*^*ERT2*^ post 32 days tamoxifen injection, D), and NMNAT1 cKO: SARM1 KO (*Nmnat1*^*fl/fl*^: *Sarm1*^*-/-*^: *ActCre*^*ERT2*^ at post 32 days tamoxifen injection, E). Blue bars represent outer nuclear layer (ONL) and inner nuclear layer (INL). Similar results were obtained from 3 mice for WT, 2 mice for NMNAT1 cKO, and 3 mice for NMNAT1cKO:SARM1 KO. (F) The quantification of the retina thickness from NMNAT1 WT and NMNAT1 cKO:SARM1 KO mice were shown. Graphs show the average and error bars represent the standard error. Statistical analysis was performed by two-way ANOVA with Tukey post-hoc test (n = 3 mice for NMNAT1 WT, n = 3 mice for NMNAT1 cKO:SARM1 dKO). F(1, 72) = 0.8, p = 0.37 between NMNAT1 WT and NMNAT1 cKO:SARM1 KO retina. There is no significant difference between NMNAT1 WT and NMNAT1 cKO:SARM1 KO. (G) The quantification of the outer nuclear layer (ONL) thickness from NMNAT1 WT and NMNAT1 cKO:SARM1 KO mice were shown. Graphs show the average and error bars represent the standard error. Statistical analysis was performed by two-way ANOVA with Tukey post-hoc test (n = 3 mice for NMNAT1 WT, n = 3 mice for NMNAT1 cKO:SARM1 dKO). F(1, 64) = 0.43, p = 0.51 between NMNAT1 WT and NMNAT1 cKO:SARM1 KO retina. There is no significant difference between NMNAT1 WT and NMNAT1 cKO:SARM1 KO. (H) Quantification of relative ONL nuclei numbers compared with WT. The graph shows all data points and median (cross bars). Statistical analysis was performed by Mann-Whitney U test (n = 3 mice for NMNAT1 WT, n = 3 mice for NMNAT1 cKO:SARM1 KO). P = 0.10. There are no statistical differences between NMNAT1 WT and NMNAT1 cKO:SARM1KO. (I, J, K) ERG analysis of NMNAT1 WT (*Nmnat1*^*fl/fl*^: *Sarm1*^*+/-*^ or *Nmnat1*^*fl/fl*^: *Sarm1*^*-/-*^), NMNAT1 cKO (*Nmnat1*^*fl/fl*^: *ActCre*^*ERT2*^ post 29 to 32 days tamoxifen injection), and NMNAT1 cKO: SARM1 KO (*Nmnat1*^*fl/fl*^: *SARM1*^*-/-*^: *ActCre*^*ERT2*^ post 32 days tamoxifen injection). Graphs show the average and error bars represent the standard error. Statistical analysis was performed by two-way ANOVA with Tukey post-hoc test (n = 8 mice for NMNAT1 WT, n = 3 mice for NMNAT1 cKO, and n = 8 mice for NMNAT1 cKO: SARM1 KO). F (2, 144) = 29, p = 2.9 × 10^−11^ among genotypes (NMNAT1 WT, NMNAT1 cKO, NMNAT1 cKO:SARM1 KO) for scotopic a-wave, F (2, 144) = 46, p < 2.0 × 10^−16^ among genotypes for scotopic b-wave, F (2, 112) = 94, p < 2.0 × 10^−16^ among genotypes for photopic b-wave. * P < 0.05, ** P < 0.001, and ***p < 0.0001 denotes the statistical difference between NMNAT1 WT and NMNAT1 cKO or between NMNAT1 cKO:SARM1KO and NMNAT1 cKO. There is no statistical difference between NMNAT1 WT and NMNAT1 cKO: SARM1 KO.

## Discussion

In this study, we demonstrate that deletion of NMNAT1 in the adult retina causes a dramatic loss of photoreceptors and a concomitant reduction in retinal function. In addition, cell-type specific deletion of NMNAT1 in early postnatal photoreceptors is sufficient to induce retinal degeneration. Hence, NMNAT1 is required for the survival and function of both developing and mature photoreceptors. Using a modified AAV8 vector and the human rhodopsin kinase promoter to express NMNAT1, we demonstrated that a gene replacement strategy can improve retinal function in this model of LCA9. Finally, we defined the molecular mechanism by which NMNAT1 promotes photoreceptor function and survival. In photoreceptors, loss of NMNAT1 leads to activation of the inducible NADase SARM1 and the SARM1-dependent degeneration of photoreceptors. This finding defines a common mechanism operant in both photoreceptor degeneration and pathological axon degeneration. Loss of NMNAT1 in photoreceptors or NMNAT2 in axons leads to the SARM1-induced death of photoreceptors or axons, respectively. This surprising result extends our understanding of both the mechanisms causing retinal degeneration and the potential role of SARM1 in human disease ^33,50^.

Retinal NAD^+^ homeostasis is crucial for visual function and NAD^+^ decline is a hallmark of many retinal degenerative disease models ^9^. Reduced NAD^+^ induces mitochondrial dysfunction in photoreceptor cells and affects activity of SIRT3, which protects the retina from light-induced and other forms of neurodegeneration. In addition, NAD^+^-dependent enzymes play crucial roles in phototransduction including the regeneration of the photosensitive element, 11-cis-retinal, and the regulation of photoreceptor membrane potential. Moreover, mutations in the genes encoding some of these enzymes cause retinal degenerative disease. For example, mutations in all-trans-retinal dehydrogenase (RDH12) that is localized to photoreceptor cells are associated with LCA13. Combined deletion of retinal dehydrogenases, RDH12 and RDH8, results in mouse retinal degeneration ^51^. NAD^+^ is also a cofactor for inosine monophosphate dehydrogenase (IMPDH1), which is the rate limiting enzyme for GTP synthesis and, in turn, is required for cGMP production. cGMP is indispensable for the regulation of photoreceptor membrane potential and calcium concentration upon light stimulation. IMPDH1 mutations cause both a dominant form of retinitis pigmentosa (RP10) and LCA11. These results highlight the central role of NAD^+^ metabolism in the photoreceptor.

NMNAT1 is the only NMNAT enzyme localized to the nucleus in mammals and is crucial for nuclear NAD^+^ synthesis. Despite the broad functions of nuclear NAD^+^ in all cell types, the sole consequence of LCA9-associated *NMNAT1* mutations is retinal dysfunction/degeneration without systemic abnormalities. Previous studies, and our results, show early loss of photoreceptor cells in NMNAT1-deficient retina ^11,12^. In photoreceptors NMNAT1 may localize not only in the nucleus but also outside the nucleus, since a subcellular proteomics study showed the existence and enrichment of NMNAT1 in the photoreceptor outer segments ^25^. Single-cell transcriptomic RNA analysis also found *Nmnat1* in rods and cones ^48,52^. Consistent with an extranuclear role for NMNAT1 in photoreceptors, in these studies cytosolic NMNAT2 was either not identified or was found at much lower levels than NMNAT1. NMNAT3 is the mitochondrial NMNAT, and we show here that it is dispensable for retinal homeostasis and function, further highlighting the central requirement for NMNAT1 in photoreceptors.

Having demonstrated that NMNAT1 is required in photoreceptors in this model of LCA9, we showed that viral mediated gene replacement in photoreceptors is capable of improving retinal function. Adeno associated virus (AAV) is a naturally occurring, non-pathogenic virus used in gene therapy studies to restore structure and function to diseased cells. Recently, the U.S. FDA approved an AAV-RPE65 vector as a therapeutic reagent for LCA2 and other biallelic RPE65 mutation associated retinal dystrophies ^53^. Theoretically, LCA9 caused by the loss of NMNAT1 function is a reasonable target for AAV mediated gene therapy. To achieve expression in photoreceptors, we used an AAV8 variant that is a highly efficient for transducing photoreceptors following subretinal injection as well as the hGRK1 promoter that has activity exclusively in rods and cones ^43,44^. Delivery of AAV8(Y733F)-hGRK1-NMNAT1 significantly improved the retinal phenotype of mice deficient for NMNAT1, however the virus was delivered prior to deletion of NMNAT1. Future studies will assess the efficacy of gene replacement after deletion of NMNAT1 to more closely mimic the human condition.

Since NAD^+^ plays such a central role in photoreceptors, the identification of the NAD^+^ biosynthetic enzyme NMNAT1 as the cause of LCA9 suggests that photoreceptor degeneration in LCA9 is due the reduction in NAD^+^ synthesis. Surprisingly, we demonstrate here that this is not the essential function for NMNAT1 in photoreceptors. Instead, NMNAT1 is required to restrain the activity of the prodegenerative NADase SARM1. When NMNAT1 is deleted from SARM1 KO photoreceptors, the photoreceptors do not die and but instead maintain their physiological function, demonstrating that these cells do not require NMNAT1 as long as SARM1 is not present. This finding is perfectly analogous to the relationship between NMNAT2 and SARM1 in the axon. NMNAT2 KO mice are perinatal lethal and have dramatic axonal defects, but NMNAT2, SARM1 double KO mice are viable and have a normal lifespan ^32^. NMNAT enzymes inhibit the activation of SARM1 ^31^, potentially by consuming the NAD^+^ precursor NMN, which is postulated to activate SARM1 ^45,46^. Prior to our current study, loss of NMNAT2 was the only known trigger of SARM1 activation. Our current work suggests that SARM1 is activated by the loss of any NMNAT enzyme whose activity is not redundant with another NMNAT isoform. NMNAT2 is the only cytosolic NMNAT in the axon, and so loss of axonal NMNAT2 leads to localized activation of SARM1 and axon degeneration. In photoreceptors, NMNAT1 is not only nuclear but also likely extranuclear, and NMNAT2 is apparently present at very low levels. Hence, in photoreceptors loss of NMNAT1 triggers activation of SARM1 which consumes NAD^+^ and triggers cell death. As NAD^+^ loss is a common pathology of many retinal diseases, this raises the possibility that SARM1 activation may contribute to a wide range of retinal disorders. In support of this conjecture, recent studies found that SARM1 promotes retinal degeneration in X-linked retinoschisis ^54^ and rhodopsin-deficient mice^55^.

Our identification of SARM1 as the executioner of photoreceptor death in this model of LCA9 opens up new therapeutic possibilities. We previously developed a potent dominant negative SARM1 variant and demonstrated that AAV-mediated expression of dominant negative SARM1 strongly protects injured axons from degeneration in the peripheral nervous system ^56^ and is also effective in a neuroinflammatory model of glaucoma ^57^. While NMNAT1 gene replacement is a potential treatment option for LCA9, if SARM1 plays a more general role in retinal degeneration, then using gene therapy to express this dominant negative SARM1 could not only treat LCA9, but also multiple retinal neurodegenerative diseases. In addition, SARM1 is an enzyme and so small molecule enzyme inhibitors would be another attractive treatment modality ^35,36^. These findings demonstrate the utility of dissecting the molecular mechanism of degeneration in diseases of retinal neurodegeneration. In the case of LCA9, these studies identified a SARM1-dependent photoreceptor cell death pathway and discovered the heretofore unknown commonality between the mechanism of retinal neurodegeneration and pathological axon degeneration.

## Supplemental figures

**Supplemental Figure1.**
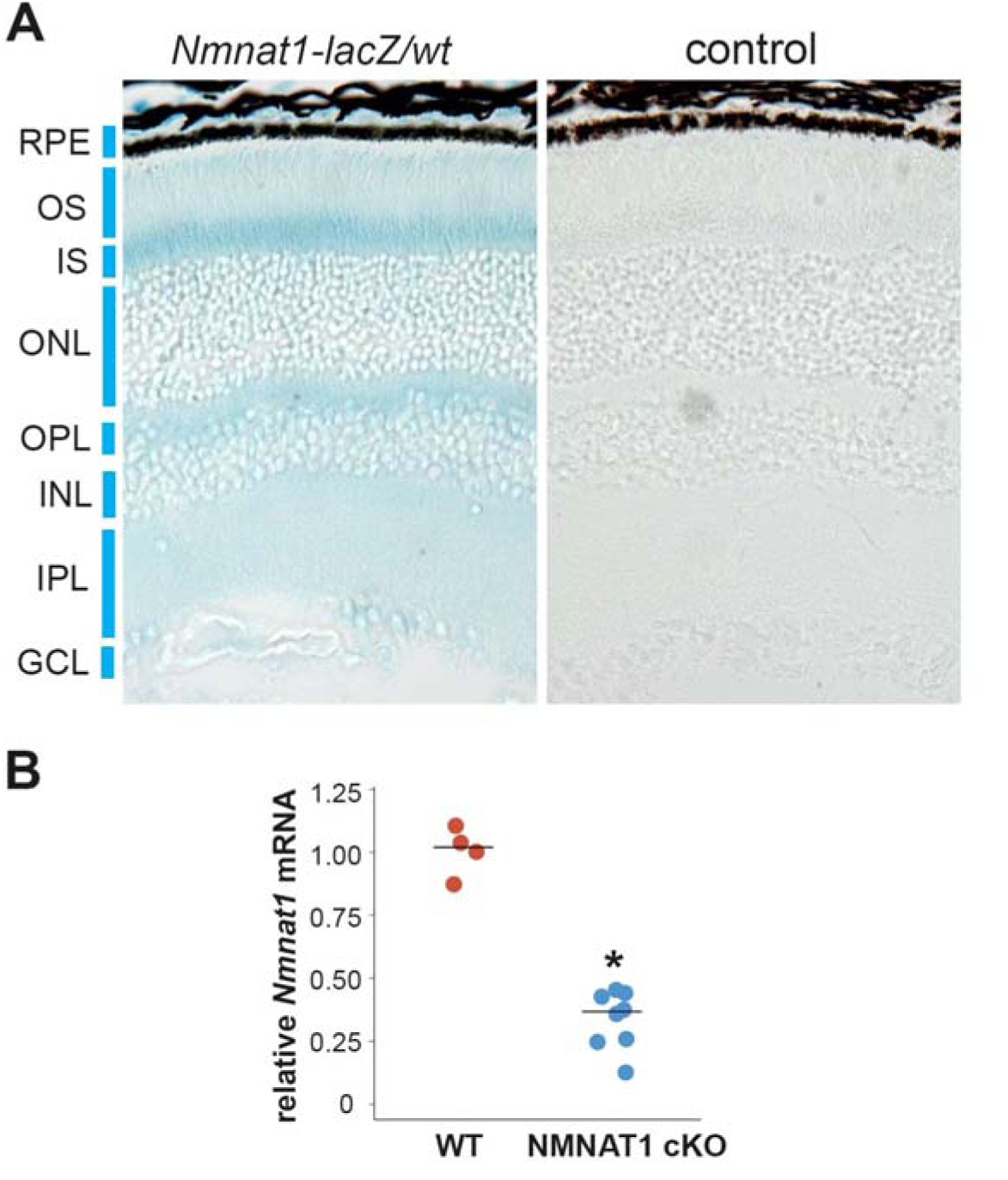
NMNAT1 is ubiquitously expressed in the retina. (A) X-Gal staining of retinal tissues from mice heterozygous for Nmnat-lacZ fusion protein lacking the nuclear localization signal (*Nmnat1-lacZ/wt*) or wild type mice (control). (B) Quantitative RT-PCR analysis of *Nmnat1* mRNA in retinal tissues from wild type (WT) or *Nmnat1*^*fl/fl*^: *ActCre*^*ERT2*^ mice at 21 days post tamoxifen injection (NMNAT1 cKO) showed significant reduction of *Nmnat1* mRNA compared with WT. * p<0.05 denotes the significant difference from WT with Mann-Whitney U test (n = 4 for WT (2 mice) and n = 8 for NMNAT1 cKO (4 mice)).

**Supplemental Figure2.**
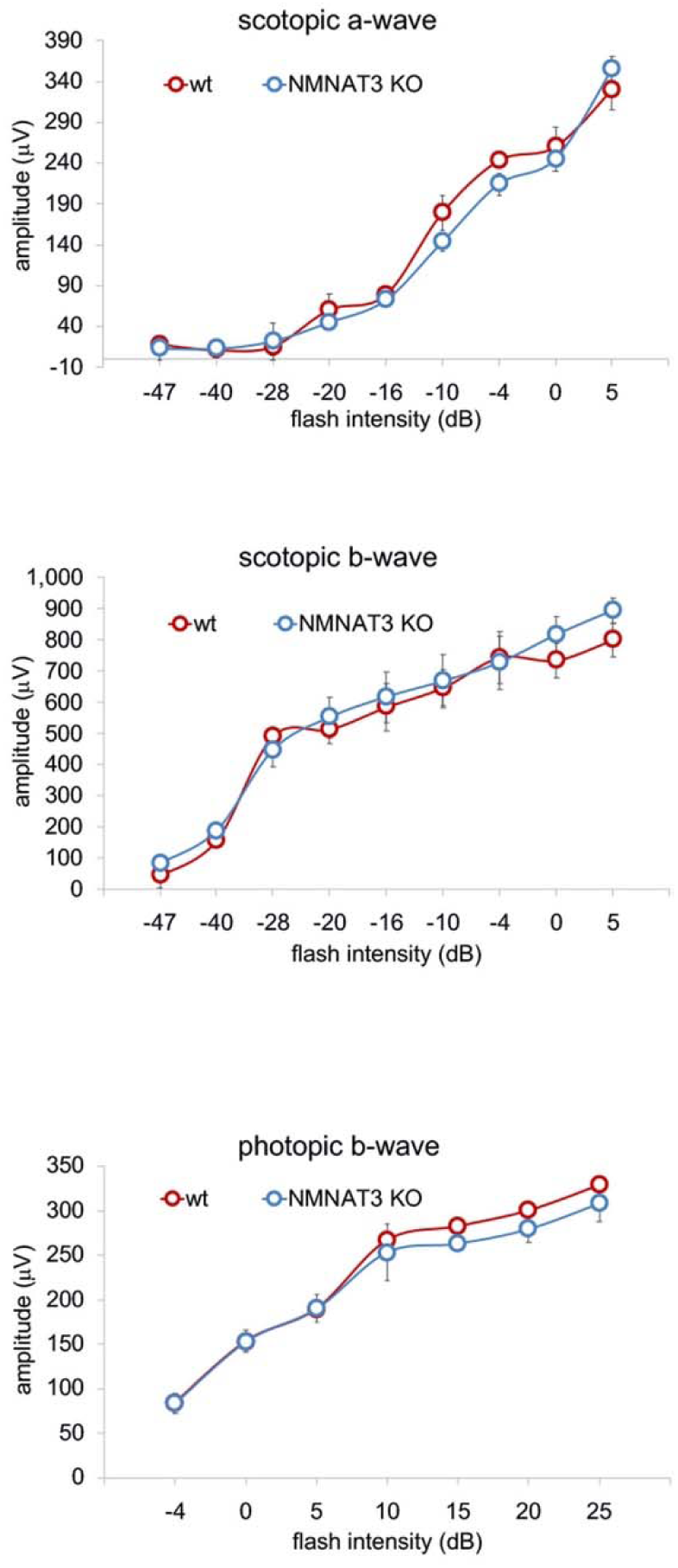
ERG analysis of NMNAT3-deficient retina. ERG analysis of WT or NMNAT3 knock out mice (NMNAT3 KO). Graphs show the average and error bars represent the standard error. Statistical analysis was performed by one-way ANOVA (n = 3 mice for WT, n = 3 mice for NMNAT3 KO). F (8, 36) = 0.78, p = 0.623 for scotopic a-wave, F (8, 36) = 0.28, p = 0.97 for scotopic b-wave, F (6, 28) = 0.23, p = 0.97 for photopic b-wave and there is no statistical difference between WT and NMNAT3 KO in each flush intensity.

**Supplemental Figure3.**
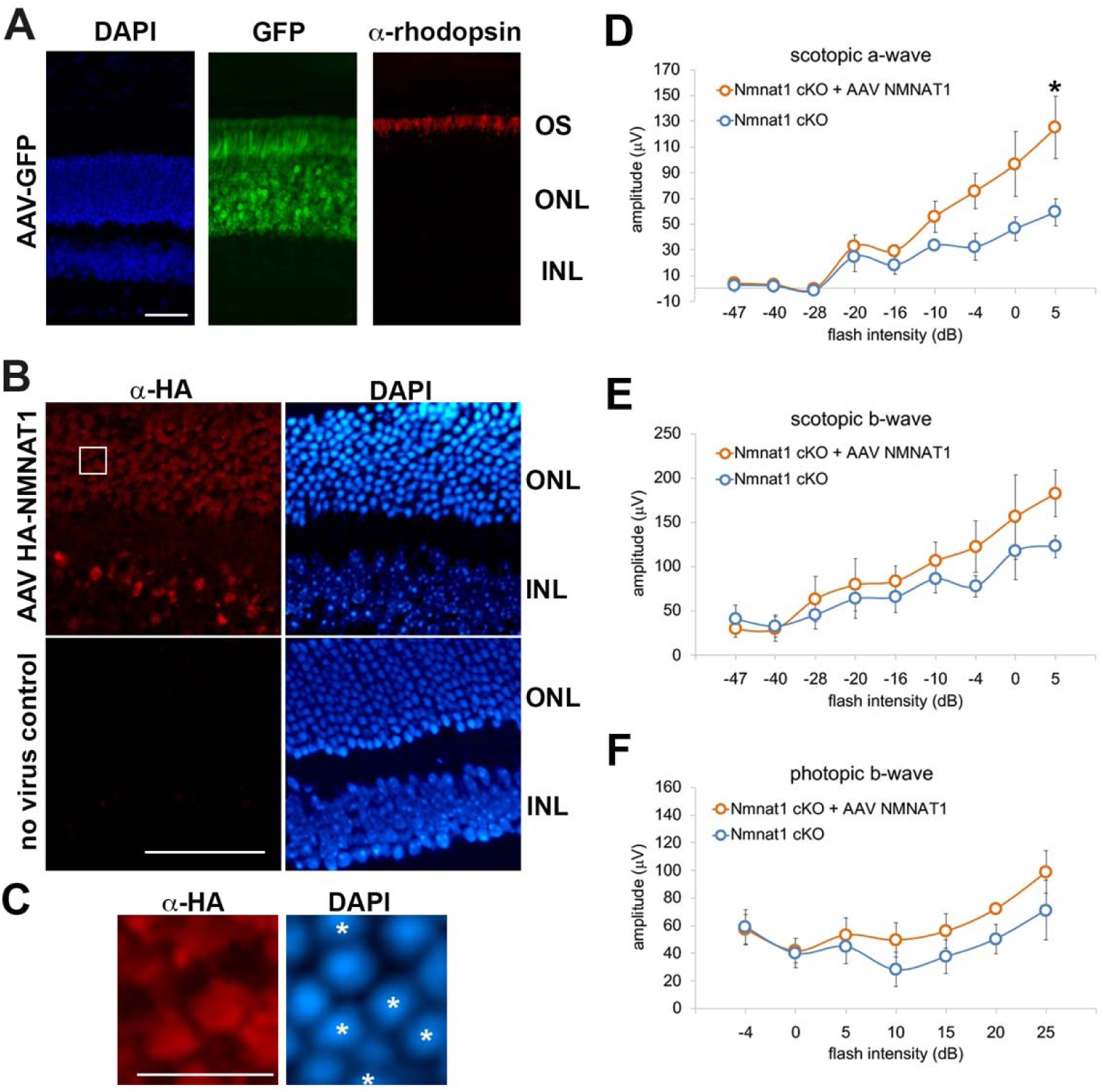
AAV-NMNAT1 partially rescued the retinal degeneration in NMNAT1-deficient retinas. (A) Fluorescent microscope images of the retina after subretinal injection of AAV expressing GFP. The photoreceptor cell layer is identified with immunostaining with antibody against human rhodopsin. Scale bar, 50 μm. (B) Fluorescent microscope images of the retina after subretinal injection of AAV expressing HA-tagged human NMNAT1. The expression of NMNAT1 is shown by immunohistochemistry with antibody against HA epitope tag and nuclei are visualized with DAPI. Asterisks indicate the cells expressing HA-Nmnat1. Scale bar, 50 μm. (C) Enlarged images corresponding to the white box in (B), showing the partial expression of NMNAT1 in the cells in the outer nuclear layer. The stars indicate the NMNAT1 expressing cells. Scale bar, 10 μm. (D, E, F) ERG analysis of AAV-GFP (control) or AAV NMNAT1 (AAV-NMNAT1) administrated NMNAT1 cKO mice. Scotopic a-wave (D), scotopic b-wave (E), and photopic b-wave amplitudes (F) are shown. Graphs show the average and error bars represent the standard error. Statistical analysis was performed by two-way ANOVA with Tukey post-hoc test (n = 5 for control or AAV-NMNAT1). F(8, 72) = 2.4, p = 0.022 between control and AAV-NMNAT1 for scotopic a-wave, F(8, 72) = 0.48, p = 0.86 between control and AAV-NMNAT1 for scotopic b-wave, F(6, 56) = 0.39, p = 0.88 between controls and AAV-NMNAT1 for photopic b-wave. *P<0.05 denotes the statistical difference between WT and AAV-NMNAT1 mice.

## Materials and methods

### Mouse

Animal studies were carried out under approved protocols from animal studies committee at Washington University. NMNAT1 mutant mice (*Nmnat1 FRTgeo;loxP/+*) which have FRT sites flanking promoterless *LacZ-neomycin phosphotransferase* gene (*beta Geo*) expression cassette located between exon 2 and 3 together with loxP sites flanking exon 3 was obtained from EUCOMM (NMNAT ^*tm1a(EUCOMM)Wtsi*^). This mouse expresses functionally null truncated NMNAT1 (exon 1 and 2) fused to beta Geo. *Nmnat1 FRTgeo;loxP*/*+* heterozygote mice were viable and fertile however, in consistent with former results, no whole body knockout (Nmnat1*FRTgeo;loxP*/*FRTgeo;loxP*) was born ^37^. Next *Nmnat1 FRTgeo;loxP*/*+* mice were crossed with FLP recombinase expressing mice in the C57BL/6 J background to remove beta Geo cassette flanked by FRT sites and RD8 mutation that might affect the ocular phenotypes ^58^. The resultant mice (*Nmnat1* ^*fl/+*^) have two loxP sites flanking the third exon. Then *Nmnat1* ^*fl/+*^ mice were crossed with mice expressing inducible Cre recombinase under actin promoter (ActCre^ERT2^) and *Nmnat1*^*fl/fl*^: *ActCre*^*ERT2*^mice were generated. All genotypes were confirmed by genomic PCR. NMNAT1 whole body knockout mice were generated by injecting 100 μg/g 4-hydroxytamoxifen (Sigma) into 6 to 8 weeks old *Nmnat1*^*fl/fl*^: ActCre^ERT2^ with IP for total 10 days with 2 days rest after first 5 days injection. The last day of injection was counted as day 0 after tamoxifen injection. Mice expressing Cre recombinase (acinCreERT2) were obtained from The Jackson Laboratory. To generate mice lacking *Nmnat1* specifically from rod photoreceptors, we crossed *Nmnat1*^*fl/fl*^ mice with mice carrying a copy of the Rhodopsin-iCre75 transgene, which were provided by Dr. Ching-Kang Jason Chen ^40^. To generate mice lacking NMNAT1 specifically from cone photoreceptors, we crossed *Nmnat1 loxP/loxP* mice with mice carrying one copy of the human red/green pigment-Cre (HGRP-Cre) transgene, which were provided by Dr. Yun Le ^42^. SARM1 knockout mice were obtained from Dr. Marco Colonna ^49^. NMNAT3 knockout mice were derived from ES cells (Nmnat3^tm1(KOMP)Mbp^, Knockout Mouse Project (KOMP)) in our facility and crossed with C57BL/6 J mice for at least five generations.

### AAV preparation

Plasmids containing the photoreceptor-specific human rhodopsin kinase (hGRK1) promoter up stream of either GFP or HA-tagged NMNAT1 were packaged in AAV8(Y733F) capsid. The detailed methodology of vector production and purification has been previously described ^59^. Briefly, vectors were packaged using a plasmid based system in HEK293 cells by CaPO_4_ transfection. Cells were harvested and lysed by successive freeze thaw cycles. Virus within the lysate was purified by discontinuous iodixanol step gradients followed by further purification via column chromatography on a 5ml HiTrap Q sepharose column using a Pharmacia AKTA FPLC system (Amersham Biosciences, Piscataway, NJ, USA). Vectors were then concentrated and buffer exchanged into Alcon BSS (Sodium-155.7 mM, Potassium-10.1 mM, Calcium-3.3 mM, Magnesium-1.5 mM, Chloride-128.9 mM, Citrate-5.8 mM, Acetate-28.6 mM, Osmolality-298 mOsm) supplemented with Tween 20 (0.014%). Virus was titered by qPCR relative to a standard and stored at −80C° as previously described ^60^.

### Subretinal Injections

Mice were anesthetized with a mixture of ketamine (70-80 mg/kg) and xylazine (15 mg/kg) injected intraperitoneally. The pupil was dilated with 1% tropicamide and topical anesthesia (0.5% proparacaine hydrochloride ophthalmic solution) was also applied to the eye. A self-sealing scleral incision was made by using the tip of a 31 G needle with the bevel pointed down. Then a 33G needle on a Hamilton syringe was inserted into the scleral incision and 1 ul of AAV containing solutions were injected in the subretinal space inducing a transient retinal detachment. The needle was slowly removed to prevent reflux and an ophthalmic ointment of neomycin/polymyxin B sulfate/bacitracin zinc was applied to the injected eye.

### Fundus microscopy and fluorescent angiography

Digital color fundus photography was performed using the Micron III retinal imaging system (Phoenix Research Laboratories). Prior to fundus imaging, mice were anesthetized with an intraperitoneal injection of 86.9 mg/kg ketamine and 10 mg/kg xylazine and administered 1.0% tropicamide eye drops (Bausch & Lomb) to dilate the pupils.

### Electroretinography (ERG)

ERG was performed as previously described (Hennig et al., 2013) by using the UTAS-E3000 Visual Electrodiagnostic System running EM for Windows (LKC Technologies). Mice were anesthetized by intra peritoneal injection of a mixture of 86.9 mg/kg ketamine and 13.4 mg/kg xylazine. The recording electrode was a platinum loop placed in a drop of methylcellulose on the surface of the cornea; a reference electrode was placed sub-dermally at the vertex of the skull and a ground electrode under the skin of the back or tail. Stimuli were brief white flashes delivered via a Ganzfeld integrating sphere, and signals were recorded with bandpass settings of 0.3 Hz to 500 Hz. After a 10-minute stabilization period, a 9-step scotopic intensity series was recorded that included rod-specific/scotopic bright flash responses. After a 5-minute light adaptation period on a steady white background, a 7-step cone-specific/photopic intensity series was recorded. Scotopic and photopic b-wave amplitudes and scotopic a-wave amplitudes were recorded for all flash intensities. We extracted quantitative measurements from the ERG waveforms using an existing Microsoft Excel macro that defines the a-wave amplitude as the difference between the average pre-trial baseline and the most negative point of the average trace and defines the b-wave amplitude as the difference between this most negative point to the highest positive point, without subtracting oscillatory potentials.

### Quantitative RT-PCR

Mice were euthanized and eyeballs were enucleated and retinas were dissected and immediately freeze in liquid N_2_. On the day of preparation, Trizol was directly added to the frozen retina and tissues were homogenized with Polytron and RNA was extracted using Trizol (Thermo Fisher Scientific) and chloroform (Sigma) phase separation. Quantitative RT-PCR reaction was performed with primers (*Nmnat1*-forward: AGAACTCACACTGGGTGGAAG, *Nmnat1*-reverse: CAGGCTTTTCCAGTGCAGGTG, Gapdh-forward: TGCCCCCATGTTTGTGATG, Gapdh-reverse: TGTGGTCATGAGCCCTTCC with reaction mixture (ThermoFisher, SYBR® Green PCR Master Mix) and monitored with Prism 7900HT (ABI) and analyzed with delta-CT method.

### Histology

Mice were euthanized and eyeballs were enucleated and fixed in 4% formalin for 8hr then washed with PBS and then embedded in paraffin. The thickness of the retinal layers or outer nuclear layers was measured using HE stained sections and plotted against the distance from the optic nerve head. The numbers of nuclei in the outer nuclear layer were analyzed using HE stained retinal sections. Outer nuclear layer was visually determined and the number of nucleus in each layer was counted and normalized by the length parallel to each layer of the retina. Data were expressed relative to the total number of nuclei in the WT. For immunostaining of the HA epitope tag, paraffin embedded eye sections were deparaffinized and treated with formic acid (70% in water) for 15min at room temperature. Sections were rinsed and treated with blocking solution (goat IgG). Primary antibody against HA (Cell Signaling Technology, 3724, 1:400) and secondary antibody Jackson Immuno Research Laboratories, AlexaFluo@568, 111-585-003) were used to visualize the HA-tagged NMNAT1. Primary antibody against rhodopsin (Abcam, ab3267, 1:500) was used to identify the photoreceptor outer segment. Slides were analyzed under the microscope (Nikon, Eclipse 80i) after the nuclear staining with DAPI and mounting (Vector Laboratories, VECTASHIELD® with DAPI, H-1200-10). For x-gal staining, retina were dissected and fixed in the cold fixation buffer (0.2% Glutaraldehyde, 5mM EGTA, 2mM MgCl2, 0.1M K-phoshate buffer pH7.2) for 1 hr, wash with detergent rinse (0.02% Igepal, 0.01% Sodium Deoxycholate, and 2mM MgCl_2_in 0.1M phosphate buffer pH 7.3), and incubated with X-Gal solution (1mg/ml X-Gal, 0.02% Igepal, 0.01% Sodium Deoxycholate, 5mM Potassium Ferricyanide, 5mM Pottassium Ferrocyanide, and 2mM MgCl_2_,0.1M phosphate buffer pH 7.3) for 10 hours in the dark at room temperature. Tissues were rinsed with PBS then fix with 4% PFA for 1 hr then paraffin sections were prepared and the sections were analyzed under the microscope (Nikon, Eclipse 80i).

### Metabolite measurement

Mice were euthanized and eyeballs were enucleated and retinas were dissected and immediately freeze in liquid N_2_. On the day of extraction, retinal tissues were homogenized in 160 μl of cold 50% MeOH solution in water using homogenizer (Branson) and then centrifuged (15,000 g, 4 °C, 10min). Clear supernatant was transferred to new tube containing 100 μl chloroform and vigorously shake then centrifuged (15,000 g, 4 °C, 10min). The chloroform extraction was repeated three times. Clear aqueous phase (120 μl) was transferred to new tube and then lyophilized and stored at −80 °C until measurement. Lyophilized samples were reconstituted with 60 μl of 5□mM ammonium formate (Sigma) and centrifuged at 12,000 x g for 10□min. Cleared supernatant was transferred to the sample tray. Serial dilutions of standards for each metabolite in 5□mM ammonium formate were used for calibration. Liquid chromatography was performed by HPLC (1290; Agilent) with Atlantis T3 (LC 2.1 × 150 mm, 3 μm; Waters) for steady-state metabolite assays ^39^. For steady-state metabolite analysis, 20 μl of samples were injected at a flow rate of 0.15□ml/min with 5□mM ammonium formate for mobile phase A and 100% methanol for mobile phase B. Metabolites were eluted with gradients of 0–10□min, 0–70% B; 10–15□min, 70% B; 16–20□min, 0% B ^39^. The metabolites were detected with a triple quadrupole mass spectrometer (6460, Agilent) under positive ESI multiple reaction monitoring (MRM) using m/z for NAD^+^:664 > 28, NMN:335 >123, cADPR: 542>428, and Nam:123 > 80. Metabolites were quantified by MassHunter quantitative analysis tool (Agilent) with standard curves and normalized by the protein amount in the sample.

### Statistical analysis

Sample number (n) was defined as a number of mice or replicates and indicated in the figure legend. Data comparisons were performed using Mann-Whitney U test, Kruskal-Wallis test, one-way ANOVA, or two-way ANOVA using R. F and P values for ANOVA were reported for each comparison in corresponding figure legends. For multiple comparisons, Holm-Bonferroni multiple comparison for one-way-ANOVA and Tukey post-hoc test for two-way-ANOVA were used.

## Acknowledgements

We thank Amy Strickland, Nina Panchenko, Kimberly Kruse, Andrea Santeford, Rachel McClarney, Simburger Kelli, Neiner Alicia, and Cassidy Menendez for technical assistance.

## References

1. Falk, M. J. et al. NMNAT1 mutations cause Leber congenital amaurosis. Nat. Genet. 44, 1040–1045 (2012).

2. Perrault, I. et al. Mutations in NMNAT1 cause Leber congenital amaurosis with early-onset severe macular and optic atrophy. Nat. Genet. 44, 975–977 (2012).

3. Koenekoop, R. K. et al. Mutations in NMNAT1 cause Leber congenital amaurosis and identify a new disease pathway for retinal degeneration. Nat. Genet. 44, 1035–1039 (2012).

4. Chiang, P.-W. et al. Exome sequencing identifies NMNAT1 mutations as a cause of Leber congenital amaurosis. Nat. Genet. 44, 972–974 (2012).

5. Coppieters, F. et al. Hidden Genetic Variation in LCA9-Associated Congenital Blindness Explained by 5’UTR Mutations and Copy-Number Variations of NMNAT1. Hum. Mutat. 36, 1188–1196 (2015).

6. Khan, A. O. et al. Genome-wide linkage and sequence analysis challenge CCDC66 as a human retinal dystrophy candidate gene and support a distinct NMNAT1-related fundus phenotype. Clin. Genet. 93, 149–154 (2018).

7. Sasaki, Y., Margolin, Z., Borgo, B., Havranek, J. J. & Milbrandt, J. Characterization of Leber Congenital Amaurosis-associated NMNAT1 Mutants. J. Biol. Chem. 290, 17228–17238 (2015).

8. Zabka, T. S. et al. Retinal toxicity, in vivo and in vitro, associated with inhibition of nicotinamide phosphoribosyltransferase. Toxicol. Sci. 144, 163–172 (2015).

9. Lin, J. B. et al. NAMPT-Mediated NAD+ Biosynthesis Is Essential for Vision In Mice. Cell Reports 17, 69–85 (2016).

10. Zhu, Y., Zhang, L., Sasaki, Y., Milbrandt, J. & Gidday, J. M. Protection of mouse retinal ganglion cell axons and soma from glaucomatous and ischemic injury by cytoplasmic overexpression of Nmnat1. Invest. Ophthalmol. Vis. Sci. 54, 25–36 (2013).

11. Eblimit, A. et al. NMNAT1 E257K variant, associated with Leber Congenital Amaurosis (LCA9), causes a mild retinal degeneration phenotype. Exp. Eye Res. 173, 32–43 (2018).

12. Wang, X., Fang, Y., Liao, R. & Wang, T. Targeted deletion of Nmnat1 in mouse retina leads to early severe retinal dystrophy. bioRxiv 210757 (2017). doi: 10.1101/210757

13. Siemiatkowska, A. M. et al. Nonpenetrance of the most frequent autosomal recessive leber congenital amaurosis mutation in NMNAT1. JAMA Ophthalmol 132, 1002–1004 (2014).

14. Greenwald, S. H. et al. Mouse Models of NMNAT1-Leber Congenital Amaurosis (LCA9) Recapitulate Key Features of the Human Disease. Am. J. Pathol. 186, 1925–1938 (2016).

15. Kuribayashi, H. et al. Roles of Nmnat1 in the survival of retinal progenitors through the regulation of pro-apoptotic gene expression via histone acetylation. Cell Death Dis 9, 891–14 (2018).

16. Zhai, R. G. et al. Drosophila NMNAT maintains neural integrity independent of its NAD synthesis activity. PLoS Biol. 4, e416 (2006).

17. Zhai, R. G. et al. NAD synthase NMNAT acts as a chaperone to protect against neurodegeneration. Nature 452, 887–891 (2008).

18. Schuster, A. et al. The phenotype of early-onset retinal degeneration in persons with RDH12 mutations. Invest. Ophthalmol. Vis. Sci. 48, 1824–1831 (2007).

19. Kennan, A. et al. Identification of an IMPDH1 mutation in autosomal dominant retinitis pigmentosa (RP10) revealed following comparative microarray analysis of transcripts derived from retinas of wild-type and Rho(-/-) mice. Hum. Mol. Genet. 11, 547–557 (2002).

20. Bowne, S. J. et al. Mutations in the inosine monophosphate dehydrogenase 1 gene (IMPDH1) cause the RP10 form of autosomal dominant retinitis pigmentosa. Hum. Mol. Genet. 11, 559–568 (2002).

21. Lin, J. B., Lin, J. B., Chen, H. C., Chen, T. & Apte, R. S. Combined SIRT3 and SIRT5 deletion is associated with inner retinal dysfunction in a mouse model of type 1 diabetes. Scientific Reports 2016 6 9, 3799–12 (2019).

22. Lin, J. B. & Apte, R. S. NAD+ and sirtuins in retinal degenerative diseases: A look at future therapies. Prog Retin Eye Res 67, 118–129 (2018).

23. Verdin, E. NAD+ in aging, metabolism, and neurodegeneration. Science 350, 1208–1213 (2015).

24. Berger, F., Lau, C., Dahlmann, M. & Ziegler, M. Subcellular compartmentation and differential catalytic properties of the three human nicotinamide mononucleotide adenylyltransferase isoforms. J. Biol. Chem. 280, 36334–36341 (2005).

25. Zhao, L. et al. Integrative subcellular proteomic analysis allows accurate prediction of human disease-causing genes. Genome Res. 26, 660–669 (2016).

26. Walker, L. J. et al. MAPK signaling promotes axonal degeneration by speeding the turnover of the axonal maintenance factor NMNAT2. Elife 6, 545 (2017).

27. Sasaki, Y., Vohra, B. P. S., Baloh, R. H. & Milbrandt, J. Transgenic mice expressing the Nmnat1 protein manifest robust delay in axonal degeneration in vivo. J. Neurosci. 29, 6526–6534 (2009).

28. Babetto, E. et al. Targeting NMNAT1 to axons and synapses transforms its neuroprotective potency in vivo. J. Neurosci. 30, 13291–13304 (2010).

29. Gilley, J. & Coleman, M. P. Endogenous Nmnat2 Is an Essential Survival Factor for Maintenance of Healthy Axons. PLoS Biol. 8, e1000300 (2010).

30. Gerdts, J., Brace, E. J., Sasaki, Y., DiAntonio, A. & Milbrandt, J. SARM1 activation triggers axon degeneration locally via NADL destruction. Science 348, 453–457 (2015).

31. Sasaki, Y., Nakagawa, T., Mao, X., DiAntonio, A. & Milbrandt, J. NMNAT1 inhibits axon degeneration via blockade of SARM1-mediated NAD+depletion. Elife 5, 1010 (2016).

32. Gilley, J., Orsomando, G., Nascimento-Ferreira, I. & Coleman, M. P. Absence of SARM1 rescues development and survival of NMNAT2-deficient axons. Cell Reports 10, 1974–1981 (2015).

33. Figley, M. D. & DiAntonio, A. The SARM1 axon degeneration pathway: control of the NAD+ metabolome regulates axon survival in health and disease. Current Opinion in Neurobiology 63, 59–66 (2020).

34. Sasaki, Y. et al. cADPR is a gene dosage-sensitive biomarker of SARM1 activity in healthy, compromised, and degenerating axons. Experimental Neurology 329, 113252 (2020).

35. DiAntonio, A. Axon degeneration: mechanistic insights lead to therapeutic opportunities for the prevention and treatment of peripheral neuropathy. Pain 160 Suppl 1, S17–S22 (2019).

36. Krauss, R., Bosanac, T., Devraj, R., Engber, T. & Hughes, R. O. Axons Matter: The Promise of Treating Neurodegenerative Disorders by Targeting SARM1-Mediated Axonal Degeneration. Trends Pharmacol. Sci. 41, 281–293 (2020).

37. Conforti, L. et al. Reducing expression of NAD+ synthesizing enzyme NMNAT1 does not affect the rate of Wallerian degeneration. FEBS J. 278, 2666–2679 (2011).

38. Slivicki, R. A., Ali, Y. O., Lu, H.-C. & Hohmann, A. G. Impact of Genetic Reduction of NMNAT2 on Chemotherapy-Induced Losses in Cell Viability In Vitro and Peripheral Neuropathy In Vivo. PLOS ONE 11, e0147620 (2016).

39. Hikosaka, K. et al. Deficiency of nicotinamide mononucleotide adenylyltransferase 3 (nmnat3) causes hemolytic anemia by altering the glycolytic flow in mature erythrocytes. J. Biol. Chem. 289, 14796–14811 (2014).

40. Li, S. et al. Rhodopsin-iCre transgenic mouse line for Cre-mediated rod-specific gene targeting. Genesis 41, 73–80 (2005).

41. Aït-Ali, N. et al. Rod-derived cone viability factor promotes cone survival by stimulating aerobic glycolysis. Cell 161, 817–832 (2015).

42. Le, Y.-Z. et al. Targeted expression of Cre recombinase to cone photoreceptors in transgenic mice. Mol. Vis. 10, 1011–1018 (2004).

43. Kay, C. N. et al. Targeting photoreceptors via intravitreal delivery using novel, capsid-mutated AAV vectors. PLOS ONE 8, e62097 (2013).

44. Boye, S. L. et al. AAV-mediated gene therapy in the guanylate cyclase (RetGC1/RetGC2) double knockout mouse model of Leber congenital amaurosis. Hum. Gene Ther. 24, 189–202 (2013).

45. Di Stefano, M. et al. A rise in NAD precursor nicotinamide mononucleotide (NMN) after injury promotes axon degeneration. Cell Death and Differentiation 2014 22:5 22, 731–742 (2015).

46. Zhao, Z. Y. et al. A Cell-Permeant Mimetic of NMN Activates SARM1 to Produce Cyclic ADP-Ribose and Induce Non-apoptotic Cell Death. iScience 15, 452–466 (2019).

47. Datta, P. et al. Accumulation of non-outer segment proteins in the outer segment underlies photoreceptor degeneration in Bardet-Biedl syndrome. Proc. Natl. Acad. Sci. U.S.A. 112, E4400–9 (2015).

48. Menon, M. et al. Single-cell transcriptomic atlas of the human retina identifies cell types associated with age-related macular degeneration. Nat Commun 10, 4902–9 (2019).

49. Szretter, K. J. et al. The immune adaptor molecule SARM modulates tumor necrosis factor alpha production and microglia activation in the brainstem and restricts West Nile Virus pathogenesis. J. Virol. 83, 9329–9338 (2009).

50. Coleman, M. P. & Höke, A. Programmed axon degeneration: from mouse to mechanism to medicine. Nature Reviews Neuroscience 2011 12:8 21, 183–196 (2020).

51. Maeda, A., Golczak, M., Maeda, T. & Palczewski, K. Limited roles of Rdh8, Rdh12, and Abca4 in all-trans-retinal clearance in mouse retina. Invest. Ophthalmol. Vis. Sci. 50, 5435–5443 (2009).

52. Lukowski, S. W. et al. A single-cell transcriptome atlas of the adult human retina. The EMBO Journal 38, e100811 (2019).

53. Apte, R. S. Gene Therapy for Retinal Degeneration. Cell 173, 5 (2018).

54. Molday, L. L., Wu, W. W. H. & Molday, R. S. Retinoschisin (RS1), the protein encoded by the X-linked retinoschisis gene, is anchored to the surface of retinal photoreceptor and bipolar cells through its interactions with a Na/K ATPase-SARM1 complex. J. Biol. Chem. 282, 32792–32801 (2007).

55. Ozaki, E. et al. SARM1 deficiency promotes rod and cone photoreceptor cell survival in a model of retinal degeneration. Life Sci Alliance 3, e201900618 (2020).

56. Geisler, S. et al. Gene therapy targeting SARM1 blocks pathological axon degeneration in mice. J. Exp. Med. 216, 294–303 (2019).

57. Ko, K. W., Milbrandt, J. & DiAntonio, A. SARM1 acts downstream of neuroinflammatory and necroptotic signaling to induce axon degeneration. The Journal of Cell Biology 219, 152 (2020).

58. Mattapallil, M. J. et al. The Rd8 mutation of the Crb1 gene is present in vendor lines of C57BL/6N mice and embryonic stem cells, and confounds ocular induced mutant phenotypes. Invest. Ophthalmol. Vis. Sci. 53, 2921–2927 (2012).

59. Zolotukhin, S. et al. Production and purification of serotype 1, 2, and 5 recombinant adeno-associated viral vectors. Methods 28, 158–167 (2002).

60. Jacobson, S. G. et al. Safety of recombinant adeno-associated virus type 2-RPE65 vector delivered by ocular subretinal injection. Mol. Ther. 13, 1074–1084 (2006).

